# Genomic Prediction of Hybrid Combinations in the Early Stages of a Maize Hybrid Breeding Pipeline

**DOI:** 10.1101/054015

**Authors:** D.C. Kadam, S.M. Potts, M.O. Bohn, A.E. Lipka, A.J. Lorenz

## Abstract

Prediction of single-cross hybrid performance has been a major goal of plant breeders since the beginning of hybrid breeding. Genomic prediction has shown to be a promising approach, but only limited studies have examined the accuracy of predicting single cross performance. Most of the studies rather focused on predicting top cross performance using single tester to determine the inbred parent’s worth in hybrid combinations. Moreover, no studies have examined the potential of predicting single crosses made among random progenies derived from a series of biparental families, which resembles the structure of germplasm comprising the initial stages of a hybrid maize breeding pipeline. The main objective of this study was to evaluate the potential of genomic prediction for identifying superior single crosses early in the breeding pipeline and optimize its application. To accomplish these objectives, we designed and analyzed a novel population of single-cross hybrids representing the Iowa Stiff Stalk Synthetic/Non-Stiff Stalk heterotic pattern commonly used in the development of North American commercial maize hybrids. The single cross prediction accuracies estimated using cross-validation ranged from 0.40 to 0.74 for grain yield, 0.68 to 0.91 for plant height and 0.54 to 0.94 for staygreen depending on the number of tested parents of the single crosses. The genomic estimated general and specific combining abilities showed a clear advantage over the use of genomic covariances among single crosses, especially when one or both parents of the single cross were untested in hybrid combinations. Overall, our results suggest that genomic prediction of the performance of single crosses made using random progenies from the early stages of the breeding pipeline holds great potential to re-design hybrid breeding and increase its efficiency.

## INTRODUCTION

Contemporary hybrid breeding programs are based on the ‘pure-line method of corn breeding’ proposed by (Shull. 1909). This method includes the development of inbreds by self-pollination followed by evaluation of selected inbreds for single-cross hybrid performance when crossed to other inbreds. A major challenge with this method is achieving adequate testing of the inbreds to evaluate performance in single cross combinations (Hallauer *et al.* 1988). In maize, heterotic groups are well defined, and production of single crosses are almost exclusively made between heterotic groups. The fullest assessment of single cross performance in maize, therefore, would be a complete factorial mating design achieved by making all between-heterotic group crosses. This would provide complete information on both general combining ability (GCA) and specific combining ability (SCA) (Comstock and Robinson. 1948). However, a full factorial among inbreds can be cost prohibitive as advanced breeding programs typically have many inbreds to evaluate, making the number of all possible single crosses extremely large. For this reason, predicting single cross performance has always been a major issue in all hybrid breeding programs (Schrag *et al.* 2009).

Several approaches have been used to evaluate the genetic merit of inbreds for single cross performance with variable success. These approaches include inbred per se performance, performance when crossed to testers (“topcross” test), best linear unbiased prediction (BLUP) using pedigrees, and molecular marker-assisted hybrid prediction. Many of these approaches have been reviewed in detail elsewhere (Schrag *et al.* 2009; Smith and Betrán. 2004). Per se performance of inbred is typically found to be a very poor predictor of single cross performance, especially for traits such as grain yield, where strong dominance effects underlie the genetic variance (Hallauer. 1977; Love and Wentz. 1914; Smith. 1986). A topcross test is an established and simple approach to assess the genetic worth of inbreds in single cross combinations (Jenkins and Brunson. 1932). However, topcross evaluation of a large number of inbreds is difficult (Albrecht *et al.* 2011) and selections based on single cross performances are performed in later stages which increases the time required for hybrid development. Bernardo. (1996a) showed that pedigree-based BLUP is useful for prediction of untested single crosses. He used pedigree-based covariance matrices among tested and untested single crosses to obtain BLUPs for untested single crosses. The correlations between observed and predicted single cross performance for prediction of crosses whose both parents have been tested in hybrid combinations were moderate (0.43-0.76). However, when one or both of the parents of the single crosses were untested, the correlations were severely decreased (Bernardo. 1996b).

The relationship between genetic distance (GD) of parental inbreds, measured by molecular markers, and heterosis has been extensively studied in maize. While it is possible to predict single cross performance using genetic distances for hybrid sets composed of both intra-and inter-heterotic pool crosses, correlations for predicting inter-heterotic group crosses only were reported to be very low (Lee *et al.* 2007; Melchinger. 1999). Two possible causes of these low prediction accuracies include (1) loose association between heterotic QTL and the molecular markers used to estimate GD and (2) opposite linkage phases between the QTL and marker alleles as generally expected with inter-heterotic hybrids (Bernardo. 1992; Charcosset *et al.* 1991). Commercial single-cross hybrids consist of only inter-heterotic group crosses, making them the only ones relevant for prediction in breeding programs. In a modified approach, prediction of hybrid performance and SCA based on only significant markers was suggested (Vuylsteke *et al.* 2000), but this approach was found to be inferior to an established GCA method. Also, extending the GCA predictions with SCA estimates from associated markers did not improve the prediction accuracy (Schrag *et al.* 2006; Schrag *et al.* 2007).

Genomic prediction is an approach that uses markers to predict the genetic value of complex traits in progeny for selection and breeding (Meuwissen *et al.* 2001). When genomic predictions are used to make selections, it is referred to as genomic selection (GS). The primary difference between GS and traditional forms of marker-assisted selection (MAS) is the simultaneous use of a large number of markers distributed genome-wide as opposed to a small set of markers linked to QTL (Heffner *et al.* 2009). Implementation of genomic prediction and selection requires the development of training populations or calibration sets consisting of individuals that have been both phenotyped and genotyped, followed by model calibration. A whole suite of genomic prediction models have been developed, each deploying different strategies to estimate genome-wide marker effects (de los Campos *et al.* 2013).

Recently, published results from simulation and experimental studies have given first indications of usefulness of genomic prediction models for hybrid performance in maize (Albrecht *et al.* 2011; Albrecht *et al.* 2014; Jacobson *et al.* 2014; Massman *et al.* 2013; Riedelsheimer *et al.* 2012; Technow *et al.* 2012; Technow *et al.* 2014; Windhausen *et al.* 2012). However, most of the experimental studies were focused on prediction of topcross performance using single tester (Albrecht et al. 2011; Albrecht et al. 2014; Jacobson et al. 2014; Riedelsheimer et al. 2012; Windhausen et al. 2012). Experimental studies on genomic prediction of single cross performance have been based on historical data consisting of established inbred parents with mixed and complex ancestry (Massman et al. 2013; Technow et al. 2014). These studies used covariances among tested and untested single crosses estimated from realized genomic relationship matrices to predict the performance of untested crosses. The prediction accuracies were high, often exceeding 0.75, even when testcross information was not available for both parents of the single cross.

Identification of superior single crosses early in the breeding pipeline would be beneficial to develop superior hybrids more quickly. The current practice of initial selection among available inbreds based on their topcross performance using single tester followed by evaluation of single crosses made among selected inbreds increases time required for the hybrid development. Moreover, not all possible hybrid combinations among available inbreds get evaluated with this approach. It is important, therefore, to study the potential of genomic prediction of single cross performance in the early stages of the breeding pipeline. With this in mind, the objective of this study was to evaluate the potential of genomic prediction for identifying superior single crosses early in the breeding pipeline. Also, we evaluated how the prediction model and the composition of the training set affected the prediction accuracy of hybrid performance. To accomplish this objective, we designed and analyzed a novel population of single-cross hybrids. The parental recombinant inbred lines (RILs) and doubled haploid lines (DHLs) were randomly selected from three Iowa Stiff Stalk Synthetic (SSS) and three Non-Stiff Stalk synthetic (NSS) biparental populations. All single-cross hybrids, therefore, represented the SSS/NSS heterotic pattern commonly used in the development of North American commercial maize hybrids. All RILs and DH lines were genotyped using genotyping by sequencing (GBS) (Elshire *et al.* 2011), which represents an affordable genotyping option that is critical to the routine use of these methods in a breeding program.

## MATERIALS AND METHODS

### Germplasm

Three SSS inbred parents (PHG39, PHJ40, and B73) and three NSS inbred parents (LH82, PHG47, and PHG84) were used for creating six bi-parental families by making each of the three possible crosses between the three SSS inbreds and also between the three NSS inbreds (Figure S1). The chosen inbred parents were identified as being both genetically diverse and superior in GCA for grain yield under high planting density (Mansfield and Mumm. 2014). A total of 217 lines were developed from crosses between these inbred parents. Approximately 10% of these lines were RILs and 90% DHLs. RILs and DHLs will be hereafter referred collectively as “inbred progenies”. The number of inbred progenies in each of the six bi-parental families ranged from 2 to 69 (Table 1). Random crosses among the inbred progenies were made between heterotic groups to produce 312 single-cross hybrids. Crosses representing each bi-parental family were balanced to the extent possible while maximizing the number of inbred progenies used in crosses (Figure 1). Completely balanced representation was not achieved due to seed limitations and comparatively fewer inbred progenies available for certain crosses. Bi-parental families are comprised of RILs or DHLs resulting from a common breeding cross (i.e., within heterotic-group parent crosses). Single crosses were grouped into nine “single-cross families”, which we defined as a group of single crosses created using parents from the same bi-parental family on each side of the heterotic pattern (Table 1). For example, a single-cross hybrid with pedigree (PHJ40×PHG39)DH-1/(PHG47×PHG84)DH-1 belongs to the same single-cross family as a single-cross hybrid with pedigree (PHJ40×PHG39)DH-2/(PHG47×PHG84)DH-2. The mean number of times an individual SSS inbred progeny was used in a cross was 6.9. The mean number of times an individual NSS inbred progeny was used in a cross was 1.8. Number of single crosses per single-cross family ranged from 19 to 51 (Table 1).

**Figure 1.**
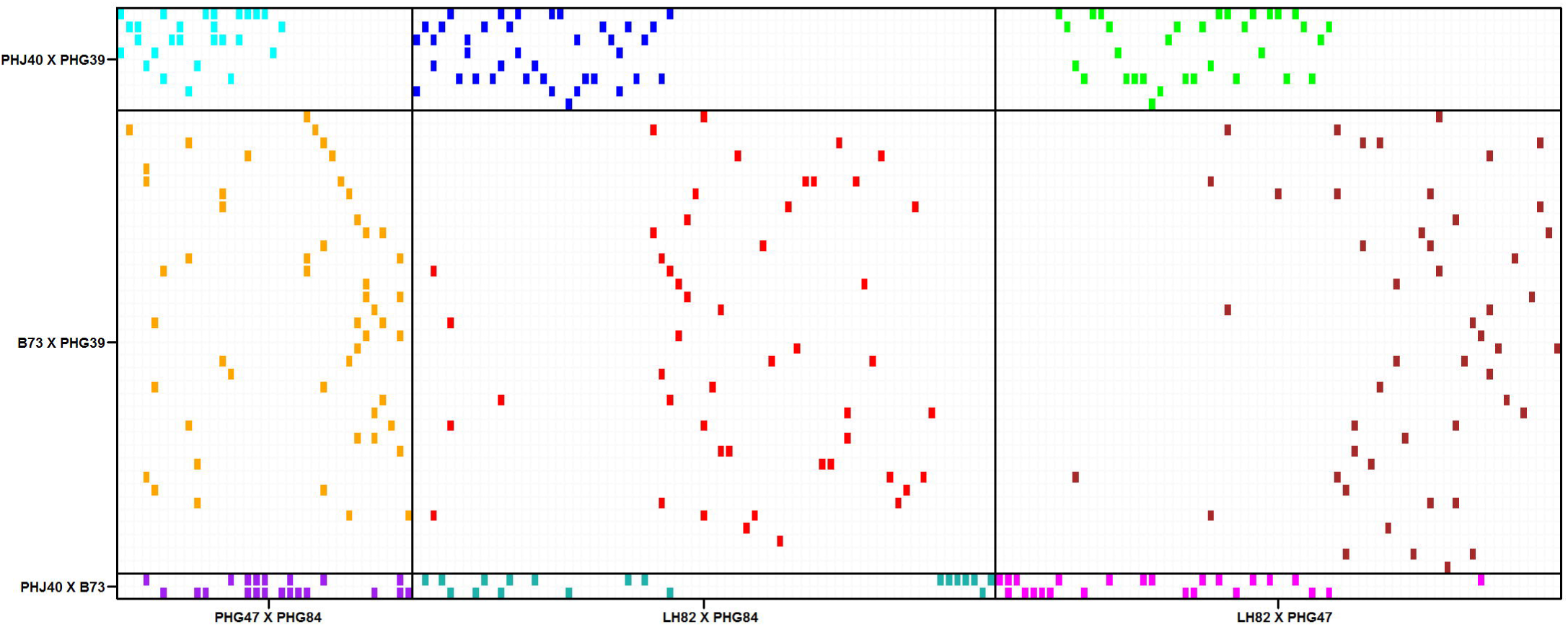
Crossing scheme between RILs or DHLs derived from three bi-parental families representing the SSS (*y*-axis) and NSS (*x*-axis) heterotic groups. Colored boxes indicate the presence while unfilled boxes indicate absence of a particular single cross. Bold lines delineate single cross families.

**Table 1.** Family designations of nine single-cross families and number of single crosses belonging to each of the nine families. Bi-parental families are listed in the row and column headings. The numbers in the parentheses indicate numbers of recombinant inbred lines (RILs) or doubled haploid lines (DHLs) in the bi-parental family or number of single crosses in each single-cross family. Total number of single crosses per bi-parental family are displayed in the table margins.

|  | PHG47xPHG84<br>(35) | LH82xPHG47<br>(69) | LH82xPHG84<br>(67) | Total |
| --- | --- | --- | --- | --- |
| PHJ40xPHG39<br>(8) | f1<br>(27) | f2<br>(39) | f3<br>(33) | **99** |
| B73xPHG39<br>(36) | f4<br>(51) | f5<br>(49) | f6<br>(49) | **149** |
| PHJ40xB73<br>(2) | f7<br>(21) | f8<br>(19) | f9<br>(24) | **64** |
| Total | **99** | **107** | **106** | **312** |

### Field experiments

The 312 single crosses were evaluated at two locations in 2012 and three locations in 2013. Two locations were common between years. The locations were as follows: South Farms (Urbana, IL; 2012 & 2013), Maxwell Farms (Urbana, IL; 2012 & 2013) and Monmouth (IL; 2013 only). The five location–year combinations were defined as separate environments. The experimental design was an α(0, 1)-incomplete block design (Patterson and Williams. 1976) with three replications at each environment. All trials were planted with an Almaco Seed Pro 360 planter set at 0.64 m row spacing and 4.46 m long row. Entries were grown in small plots consisting of two rows. Plots were overplanted by 15% to compensate for germination failure and later thinned to the target plant density of 116,000 plants ha^−1^. All fields were controlled for weeds. Nitrogen (N) was applied before planting as 28% urea-ammonium nitrate at a rate of 336.4 kg ha^−1^ to all fields. Phosphorous and potassium were each applied at 112 kg ha^−1^ according to recommended levels determined by soil tests performed by the University of Illinois Crop Science Research and Education Center. Stand counts were recorded and plots with planting densities lower than 106,000 plants ha^−1^ discarded. Additionally, issues with seed production resulted fewer hybrids being planted at all locations in 2013 (South Farms: 260; Maxwell Farms: 259 & Monmouth: 258). Plots were machine harvested and data were recorded for grain yield and several other agronomic traits. For this study, data on grain yield (GY), plant height (PH) and staygreen (SG) were used for downstream analyses. Grain yield was converted to Mt ha^−1^ on a 155 g kg^−1^ moisture basis. Plant height was measured post anthesis on a single representative plant determined by visually surveying the entire plot before measurement. Staygreen was evaluated visually as a percentage of total dry down, where a rating of 1 represented complete senescence and a rating of 10 represented fully green leaves.

### Genotyping by sequencing

Five plants of each RIL or DH were germinated. A total of 0.1 g of tissue was sampled from leaf tips and pooled across the five plants. DNA was extracted using the Qiagen DNeasy Plant 96 kit following the DNeasy Plant Handbook. DNA samples were sent to the Institute for Genomic Diversity (IGD) at Cornell University for genotyping by sequencing (GBS) where library construction and sequencing was performed as described by (Elshire *et al.* 2011). Single nucleotide polymorphisms (SNPs) were scored from the raw sequence data using the TASSEL GBS Pipeline version 3.0 (Glaubitz *et al.* 2014). SNPs with greater than 20% missing values and less than 5% minor-allele frequency were removed from the dataset. Heterozygotes were treated as missing data. Missing data was imputed using naïve imputation. Of the markers remaining after filtration, markers that were polymorphic among both SSS and NSS progenies were retained for analysis. The final marker data set consisted of 2296 high-quality SNPs. The distribution of SNPs among the 10 chromosomes of the B73 reference genome is displayed in Figure S2.

### Phenotypic data analysis

The phenotypic data were unbalanced due to missing observations. We used the following statistical model for the analysis of the data across the five environments

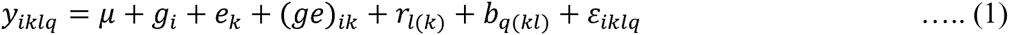

where *y*_*iklq*_ is the phenotypic observation for *i*^th^ hybrid evaluated in the *k*^th^ environment in the *l*^th^ complete block (i.e. replicate) and *q*^th^ incomplete block. The effects in the model are as follows: *μ* is the grand mean; *g*_*i*_ represents effect of the *i*^th^ hybrid; *e*_*k*_ represents the effect of the *k*^th^ environment; (*ge*)_*ik*_ represents the interaction effect between hybrid and environment; *r*_*l*(*k*)_ represents the effect of the *l*^th^ complete block nested within the *k*^th^ environment; *b*_*q*(*kl*)_ represents the effect of the *q*^th^ incomplete block nested within the *l*^th^ complete block in the *k*^th^ environment; and *ε*_*iklq*_ represents the residual. Environment and replication nested within environment effects were modeled as fixed effects while all other effects were treated as random. The distribution of *g*_*i*_ was as follow: *g*_*i*_ ~ *N*(0, *Iσ*^2^). Error and block variances were allowed to be heterogeneous among environments.

The above model was implemented using ASReml-R software (Butler *et al*. 2009) to obtain restricted maximum likelihood estimates (REML) of all variance components and solve the mixed linear model equations. Significance of the variance components was determined using likelihood ratio tests at 0.001 level of significance. The entry-mean heritability of each trait was computed according to (Holland *et al.* 2003) as: 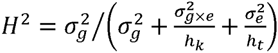, where, 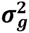 represents the variance among hybrids, 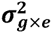 represents the variance of interaction effects of hybrids with environments, 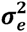 is the residual variance, *h*_*k*_ is the harmonic mean of number of observations per hybrid within an environment, and *h*_*t*_ is the harmonic mean of total number of observations per hybrid. Similarly, model (1) used to estimate the genetic variance and broad sense heritability for individual single-cross family. Finally, we calculated best linear unbiased predictions (BLUP) of hybrids and used these to evaluate hybrid prediction accuracy in further analyses.

### Genomic hybrid prediction model

We used genomic best linear unbiased prediction (G-BLUP) of untested hybrid performance. The G-BLUP model was as below:

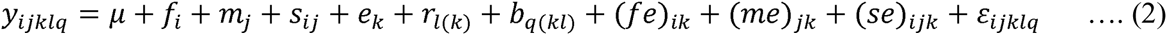

where *y*_*ijklq*_ is the phenotypic observation of a single-cross hybrid between the *i*^th^ and *j*^th^ inbred progeny evaluated in the *k*^th^ environment in the *l*^th^ complete block and *q*^th^ incomplete block. The effects in the model are as follows: *μ* is the grand mean; *f*_*i*_ and *m*_*j*_ represents the GCA effects of the female (SSS inbreds) and males (NSS inbreds), respectively; *s*_*ij*_ represents the SCA effect of the single cross; (*fe*)_*ik*_, (*me*)_*jk*_, *and* (*se*)_*ijk*_ represent the interaction effects of respective terms with the k^th^ environment. The remaining terms are as described in the model (1).

The random effect vectors *f*, *m*, and *s* were assumed to have the following multivariate normal distributions: 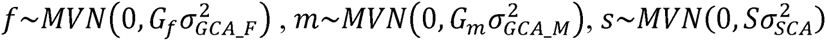 where *G*_*f*_ and *G*_*m*_ were additive genomic relationship matrices of females and males, respectively, calculated according to Method 1 of (VanRaden. 2008). The dominance relationship matrix, *S*, was computed according to (Bernardo. 2002) using the corresponding elements from matrices *G*_*f*_ and *G*_*m*_. The above model (2) was implemented using ASReml-R software (Butler et al. 2009).

We evaluated four methods to predict single cross performance using the above G-BLUP model. Broadly, these methods can be grouped into two categories: 1. Parent GCA and SCA effects; 2. Additive and dominance covariances among single crosses.

#### 1a. Parent GCA

Performance of untested single crosses (*ŷ*_*u*_) was predicted from the GCA of the corresponding parents, *i* and *j* estimated from model (2) as

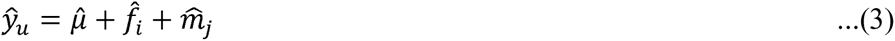

GCA of female and male lines with no performance data of their hybrids were estimated from related inbred progenies using the additive genomic relationship matrix in the linear mixed model analysis.

#### 1b. Parent GCA plus single-cross SCA

Performance of untested single crosses (*ŷ*_*u*_) was predicted using the sum of parent GCA and SCA of the crosses as

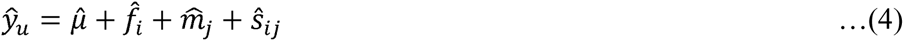
Like the GCA effects, the SCA effects for untested crosses were estimated using the dominance genomic relationship matrix in the linear mixed model analysis from model (2).

#### 2a. Additive genetic covariance among single crosses

The performance of untested single crosses (*ŷ*_*u*_) was predicted based on the covariance among tested and untested single crosses as

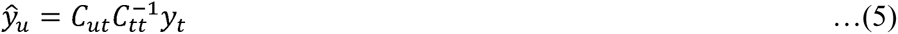

Where, *C*_*ut*_ is the genetic covariance matrix of untested and tested single crosses, *C*_*tt*_ is the phenotypic covariance matrix of the tested single crosses and *y*_*t*_ is a vector of single cross BLUPs obtained from model (1). The elements of *C*_*ut*_ and *C*_*tt*_ were computed according to (Bernardo. 2002) using the genomic relationship matrices *G*_*f*_ and *G*_*m*_. Briefly, let *i* and *i*′ denote any two female inbred progenies and *j* and *j*′ any two male inbred progenies. For a given pair of single crosses, (*i* × *j*) and (*i*′ × *j*′), the elements of *C*_*ut*_ and the off diagonal elements of *C*_*tt*_ were calculated as 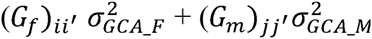. The diagonal elements of *C*_*tt*_ were estimated as 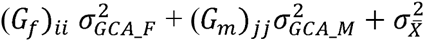 where 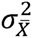 was equal to 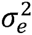 divided by the number of observations for single cross(*i* × *j*). The estimates of 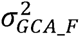 and 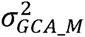 were obtained from model (2).

#### 2b. Additive plus dominance covariance among single crosses

The method described in 2a was extended by including dominance covariance among the tested and untested hybrids. Specifically, the elements of *C*_*ut*_ and off diagonal elements of *C*_*tt*_ were computed as 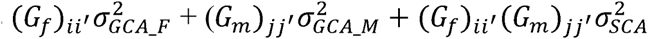. The diagonal elements of *C*_*tt*_ were estimated as 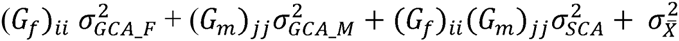. The estimates of 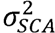 was obtained from model (2).

### Cross-validation and prediction accuracy

Accuracy of single-cross hybrid prediction was evaluated using leave-one-individual-out and leave-one-family-out cross-validations. In case of leave-one-individual-out cross-validation, the set of hybrids was divided into two subsets. The training set comprised *n*–1 hybrids (tested hybrids) and the remaining one hybrid (untested hybrid) formed the test set. This procedure was repeated *n* times such that each hybrid was placed into the “untested” set. Four scenarios involving varying degrees of tested and untested hybrids were considered. Hybrid types are those having both (T2), either female (T1F) or male (T1M), or no (T0) parental inbred evaluated for their performance in hybrid combination (Figure 2). For each cross-validation run, phenotypic data of the training set were analyzed separately and variance components were re-estimated.

**Figure 2.**
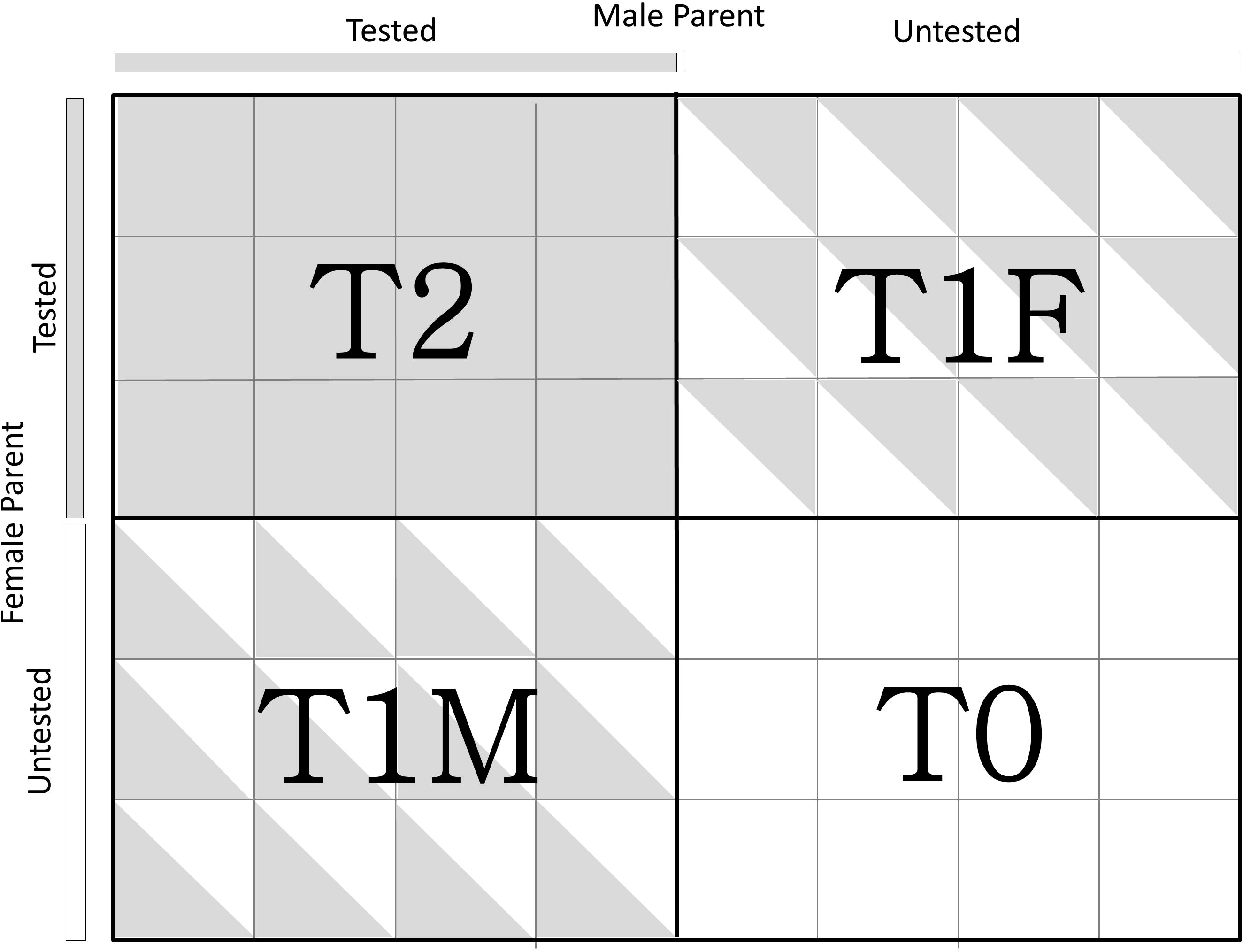
Schematic visualization of T2, T1F, T1M and T0 cross-validation scenarios.

The performance of the hybrid comprising the test set is then predicted using the four methods as described above. For leave-one-family-out cross-validations, hybrids from one of the nine families constituted the test set and remaining eight families were included in the training set. Only six of the nine single-cross families were used as test set families because of small size of the remaining three families. For both leave-one-individual-out and leave-one-family-out cross-validations, training set size was fixed to 250 in order to compare the single cross prediction accuracy of all cross-validations under a common training set size. The sampling of training sets comprised of 250 single crosses out of the single crosses from the whole set was replicated 30 times.

The hybrid BLUPs estimated from the phenotypic data were treated as the observed hybrid performance and used as the basis to evaluate hybrid prediction methods. Prediction accuracy was expressed as the Pearson’s correlation coefficient between the observed and predicted hybrid performance divided by the square root of the broad-sense heritability on an entry-mean basis (Dekkers. 2007). The mean prediction accuracy across the 30 samples of training set (n = 250) was reported. Standard errors of the prediction accuracy were calculated using the bootstrap method implemented in the R package *boot* (Canty. 2014). Briefly, for each cross-validation run, the predicted and observed values were resampled with replacement for 200 times. The distribution of 200 correlation coefficient estimates was used to estimate the bootstrap SE. This procedure was repeated for each of 30 replicates of the cross validations and the mean standard error of 30 cross validation replications was reported.

## RESULTS

### Variance components and broad-sense heritability

Variance among single-cross hybrids 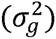 was significantly different from zero (*α* = 0.001) in the entire population as well as within single cross families for all the three traits (Table 2). For GY, the entry-mean heritability was 0.58 across the population of entire single crosses and it ranged from 0.53 to 0.83 within individual single cross families. Similarly, for PH and SG, the entry-mean heritability was 0.89 and 0.81 in the whole population, respectively, and ranged from 0.88 to 0.91 and 0.67 to 0.80 within individual single cross families, respectively. The sum of parent 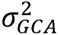 was greater than 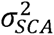 for all traits. The proportion of 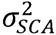 was highest for GY, followed by PH and SG (Table 3).

**Table 2.** Mean, range, genetic variance and broad-sense heritability estimates in whole population as well as individual single-cross families for grain yield (GY; Mt/ha), plant height (PH; cm), and staygreen (SG; 1-10 rating).

| Trait | Statistic | Single-cross Populations |
| --- | --- | --- |

individual single-cross families for grain yield (GY; Mt/ha), plant height (PH; cm), and staygreen (SG; 1-10 rating).
|  |  | Whole | f1 | f2 | f3 | f4 | f5 | f6 |
| --- | --- | --- | --- | --- | --- | --- | --- | --- |
| GY | Mean | 8.67 | 8.6 | 8.85 | 8.87 | 8.88 | 9.03 | 9.13 |
| (Mt ha<sup>-1</sup>) | Range | 7.14-10.2 | 7.13-9.91 | 6.94-9.94 | 7.79-9.99 | 6.74-10.7 | 7.52-10.4 | 7.46-10.5 |
| | $\sigma_g^2 \pm SE$ | $0.50 \pm 0.07$ | $0.9 \pm 0.31$ | $0.48 \pm 0.18$ | $0.25 \pm 0.12$ | $0.55 \pm 0.19$ | $0.51 \pm 0.15$ | $0.51 \pm 0.16$ |
| | $H^2 \pm SE$ | $0.58 \pm 0.04$ | $0.80 \pm 0.07$ | $0.66 \pm 0.09$ | $0.53 \pm 0.13$ | $0.57 \pm 0.10$ | $0.71 \pm 0.07$ | $0.71 \pm 0.07$ |
| PH | Mean | 210.1 | 213.4 | 206.6 | 205.7 | 221.2 | 208.9 | 216.1 |
| (cm) | Range | 191 - 231 | 197 - 227 | 187-222 | 187-222 | 202-243 | 182-230 | 191 - 241 |
| | $\sigma_g^2 \pm SE$ | $1.18 \pm 0.1$ | $0.71 \pm 0.23$ | $0.8 \pm 0.22$ | $0.95 \pm 0.27$ | $0.87 \pm 0.21$ | $0.9 \pm 0.20$ | $1.07 \pm 0.24$ |
| | $H^2 \pm SE$ | $0.89 \pm 0.01$ | $0.88 \pm 0.04$ | $0.86 \pm 0.04$ | $0.90 \pm 0.03$ | $0.83 \pm 0.04$ | $0.90 \pm 0.02$ | $0.91 \pm 0.02$ |
| SG | Mean | 6.79 | 6.96 | 7.05 | 6.68 | 6.35 | 6.75 | 6.22 |
| (1-10 rating) | Range | 5.48-8.31 | 5.57-7.96 | 5.82-8.5 | 5.57-7.96 | 4.61-7.88 | 5.67-7.92 | 5.07-7.39 |
| | $\sigma_g^2 \pm SE$ | $0.69 \pm 0.07$ | $0.36 \pm 0.15$ | $0.52 \pm 0.16$ | $0.26 \pm 0.1$ | $0.58 \pm 0.14$ | $0.38 \pm 0.1$ | $0.26 \pm 0.07$ |
| | $H^2 \pm SE$ | $0.81 \pm 0.02$ | $0.67 \pm 0.10$ | $0.74 \pm 0.07$ | $0.68 \pm 0.09$ | $0.80 \pm 0.04$ | $0.78 \pm 0.05$ | $0.78 \pm 0.05$ |

**Table 3.**
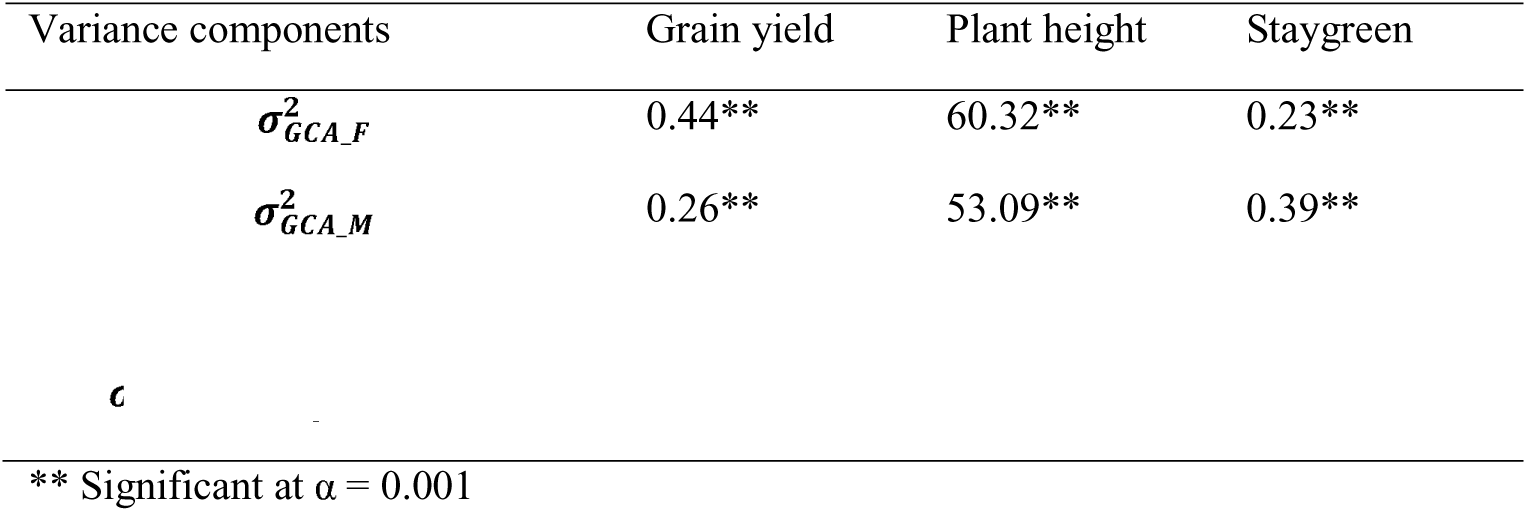
General combining ability variance of stiff stalk synthetic 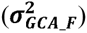 and non-stiff stalk 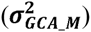 inbred progenies and specific combining ability variance 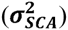 of single crosses between them.

| Variance components | Grain yield | Plant height | Staygreen |
| --- | --- | --- | --- |
| $\sigma_{GCA_F}^2$ | 0.44** | 60.32** | 0.23** |
| $\sigma_{GCA_M}^2$ | 0.26** | 53.09** | 0.39** |

\*\* Significant at $\alpha = 0.001$

### Prediction accuracy for T2, T1F, T1M and T0 scenarios

We first evaluated the prediction accuracy for T2, T1F, T1M and T0 scenarios in the entire population using leave-one-individual-out cross-validation. Higher prediction accuracies were observed for SG and PH compared to GY for all scenarios (Figure 3). Prediction accuracies were highest for T2, followed by T1F, T1M and T0. The four methods were similar in accuracy when applied to the T2 and T1F cross-validation scenarios. However, methods 1a and 1b were mostly better than methods 2a and 2b for predicting single-cross hybrid performance in the T1M and T0 scenarios. Modelling SCA led to small increases in prediction accuracy for GY and PH with a maximum increase under the T0 scenario (Table S1).

**Figure 3.**
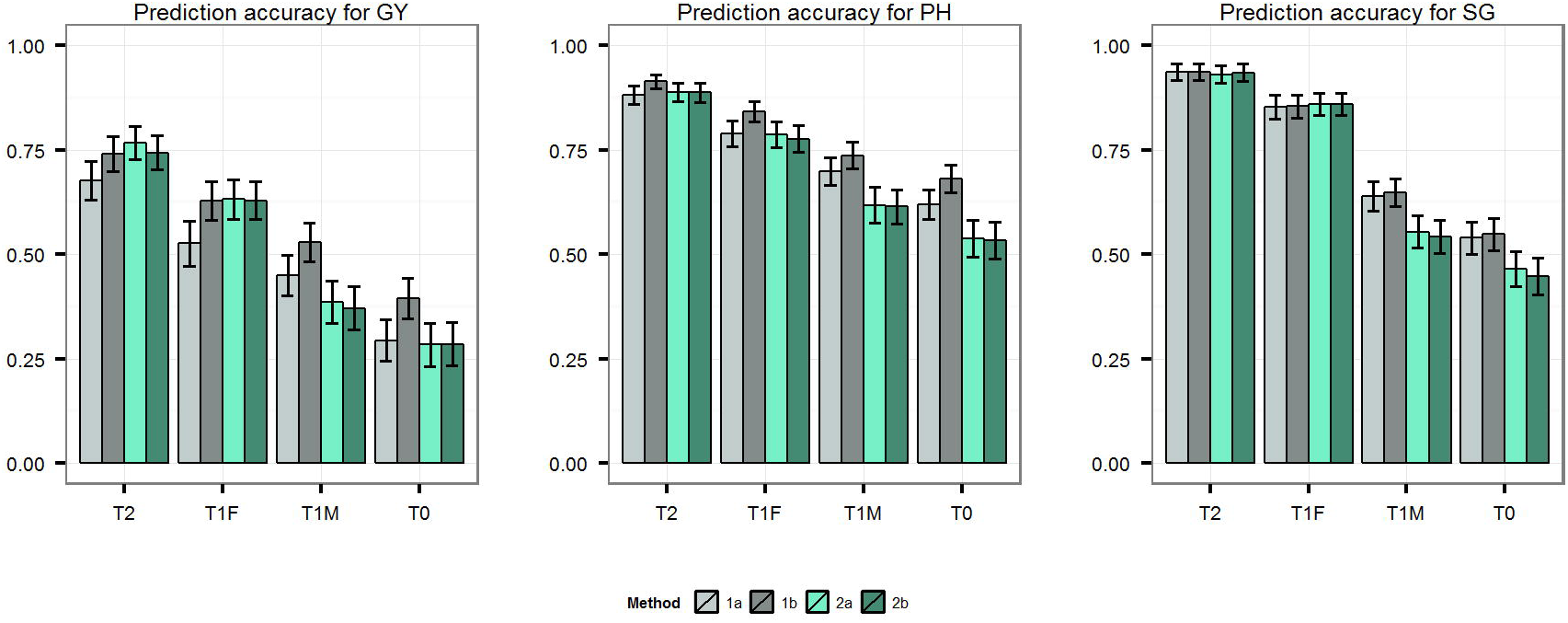
Prediction accuracies estimated using leave-one-individual-out cross validation for each genomic prediction method and cross validation scheme. Traits analyzed were grain yield (GY), plant height (PH), and stay green (SG).

### Prediction accuracy for novel family

We next investigated the potential to predict the performance of hybrids in a new single-cross family using the phenotypic and genotypic information on the hybrids from related single-cross families. The family size of three single-cross families was too small to obtain robust correlation coefficient estimates (Table 1). Hence, only six of the nine single-cross families were used as test set families. When eight of the families were used as training set to predict hybrid performances within the remaining family, prediction accuracies were generally moderate for GY and high for PH and SG (Figure 4). The mean within family accuracies were with methods 1a and 1b were 0.67, 0.62 for GY, 0.85, 0.76 for PH and 0.78, 0.78 for SG respectively (Table S2). Some variation in prediction accuracy across families was observed, especially for GY. We also evaluated the effect of adding hybrids from the family being predicted to the training set by comparing prediction accuracy of individual family with leave-one-individual-out and leave-one-family-out cross-validations. The goal of this analysis was to measure the benefit of including information from the same single-cross family to accurately separate single crosses from the same family. Although the prediction accuracies were increased slightly for some families, they were decreased for other families, indicating variable effects across families (Figure 4). The mean prediction accuracies with method 1a and 1b were 0.603, 606 for GY, 0.838, 0.85 for PH and 0.788, 0.783 for SG respectively. Method 1a and 1b showed similar prediction accuracies under both leave-one-individual-out and leave-one-family-out cross-validations for PH and SG. However, in case of GY, highest prediction accuracy was obtained by different methods for different families. Adding individuals from the family being predicted (leave-one-individual-out cross-validations) seemed to benefit method 1b more than method (Table S2).

**Figure 4.**
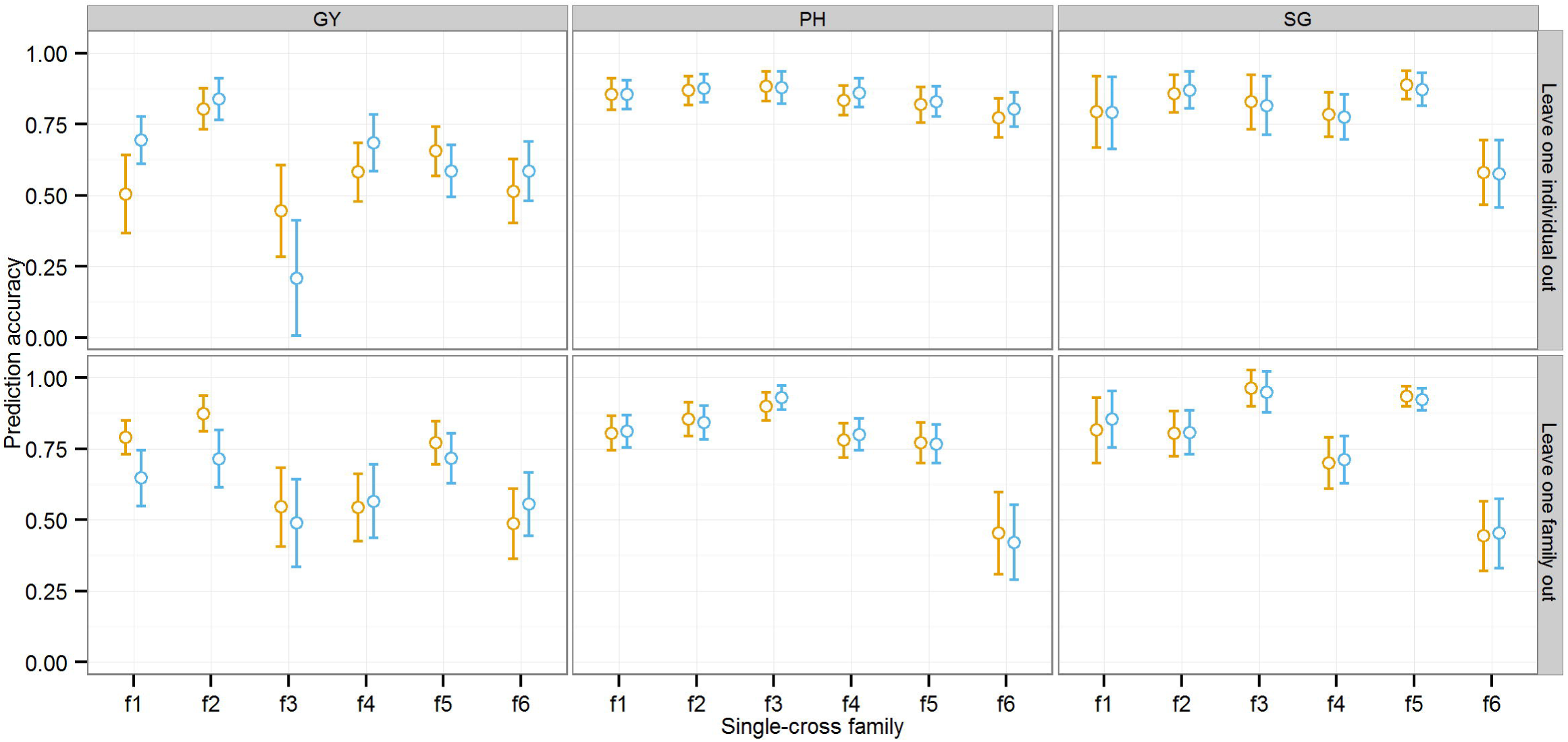
Mean prediction accuracy and standard errors of methods 1a (orange) and 1b (blue) in predicting performance of hybrids within single-cross families. Two cross-validation schemes were used: leave-one-family out (bottom panel) and leave-one-individual out (top panel). Traits analyzed were grain yield (GY), plant height (PH), and stay green (SG). Standard errors were estimated using the bootstrap method.

### Genomic predictions of grain yield of all possible single crosses

Genomic predictions were calculated for all possible 7866 single crosses between 46 SSS and 171 NSS inbred progenies based on the prediction model including parent GCA and cross SCA effects (i.e., Method 1b). The genomic predictions for GY ranged from 7590-9515 kg ha^−1^. The top 100 crosses based on genomic predictions included only one cross that was actually made and tested; the remaining 99 crosses were never made. Moreover, more than 50 untested single-cross combinations surpassed the highest genomic prediction of any tested hybrid (Figure 5).

**Figure 5.**
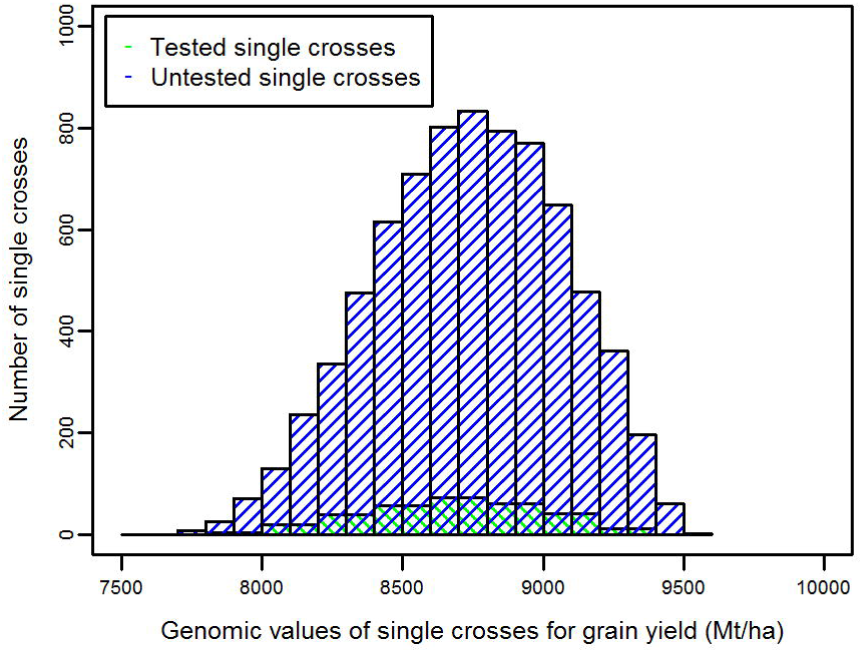
Distribution of genomic predictions for all 7866 possible single crosses between the 46 SSS inbred progenies and 171 NSS inbred progenies.

## DISCUSSION

Typical hybrid maize breeding programs involve the creation of large bi-parental families for topcrossing to elite testers early in the breeding pipeline. Early-stage selections are performed on the basis of topcross performance with single elite tester, which is the sum of the candidate line GCA effect and any SCA effect between the candidate line and elite tester used for topcrossing. While this is a very convenient and routine method, it is recognized that it would be ideal to test all combinations of possible parents immediately in the hybrid breeding pipeline (Fehr. 1987). There are two main advantages of early evaluation of all potential single crosses. First, it would help identify the best single-cross hybrid without uncertainty. If inbred progenies are selected only on the basis of topcross evaluation, there remains the possibility that some unique parental combinations never made and evaluated could actually be commercially superior products (Bernardo. 2002). Secondly, early evaluation based on single cross performance would enable the development of hybrids in shorter duration of time by essentially skipping the topcross test and immediately going to single cross evaluation. Despite these advantages, field testing of all potential single crosses of inbred progenies is completely impractical for a mature hybrid maize breeding program.

Advances in genotyping technology, such as GBS, has made it very practical to genotype all parental candidate lines with dense, genome-wide markers (He *et al.* 2014). Genomic prediction models can predict the performance of all possible single cross combinations, allowing the in-silico evaluation of all parental combinations just as in the ideal scenario. In the present study, GBS and yield trial data was used to build genomic prediction models for predicting single cross performance. The single cross prediction accuracies estimated using cross-validation ranged from 0.40 to 0.74 for grain yield, 0.68 to 0.91 for plant height and 0.54 to 0.94 for staygreen depending on the number of tested parents of the single crosses. These prediction accuracies were 52-97, 72-96 and 60-100 percent of phenotype based accuracy 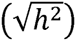 for GY, PH and SG respectively. The prediction accuracies of single cross performance achieved in this study, therefore, indicate that this approach holds great potential for increasing the efficiency of a hybrid breeding program by enabling the effective evaluation of all parental combinations.

### Prediction accuracy for T2, T1 and T0 hybrids

In order to understand the effect of tested versus untested parental lines, we evaluated the accuracies of prediction of hybrids having both (T2), either male or female (T1F and T1M), or no (T0) parental lines evaluated for their hybrid performance. The differences in prediction accuracies of T2, T1 and T0 hybrids were considerable, with the highest prediction accuracy for T2 hybrids followed by T1 hybrids and T0 hybrids. The T0 scenario was the most difficult to predict. Similar trends have been observed using simulations (Technow *et al.* 2012) as well as experimental studies based on historical data in maize (Massman *et al.* 2013; Technow *et al.* 2014), and wheat (Zhao *et al.* 2015). This finding can be explained by the representation of parents among a differing number of hybrid combinations in the training set. As the number of hybrid combinations for each parent increases, the information shared between the single crosses being predicted and the training set increases. As a result, the GCA and SCA effects are estimated with high accuracy as indicated by decreases in the standard errors with higher the number of training set hybrids parent involved (Figure 6). In the T2 scenario, both parents are represented in multiple hybrid combinations within the training set, enabling accurate estimation of parent GCA effects. With a preponderance of GCA variance over SCA variance, genotypic values of T2 hybrids can, therefore, be predicted with higher accuracy. In the case of T1 hybrids, however, only one of the parents is represented in hybrid combination. Consequently, the prediction accuracy of T1 hybrids is lower than for T2 hybrids. In the present study, the prediction accuracy of the T1F hybrids is greater than that of the T1M hybrids. This finding can be explained by the smaller total number of female than male lines, which increases the number of times each female occures in hybrid combinations. Nevertheless, the mean of T1 hybrid prediction accuracies were 78, 86 and 80 percent of the T2 hybrid prediction for GY, PH, and SG, respectively. The mean T0 hybrid prediction accuracies were 53, 75 and 59 percent of the T2 hybrid prediction accuracy for GY, PH and SG, respectively. This indicates that performance of hybrids having one untested parent can be effectively predicted using genomic estimated GCA and SCA effects, but prediction accuracies suffer considerably more if both parents are untested. This issue should be studied using larger population sizes – both in terms of more interconnected bi-parental populations and progenies per population – to determine if population size can overcome parent representation in the training set.

**Figure 6.**
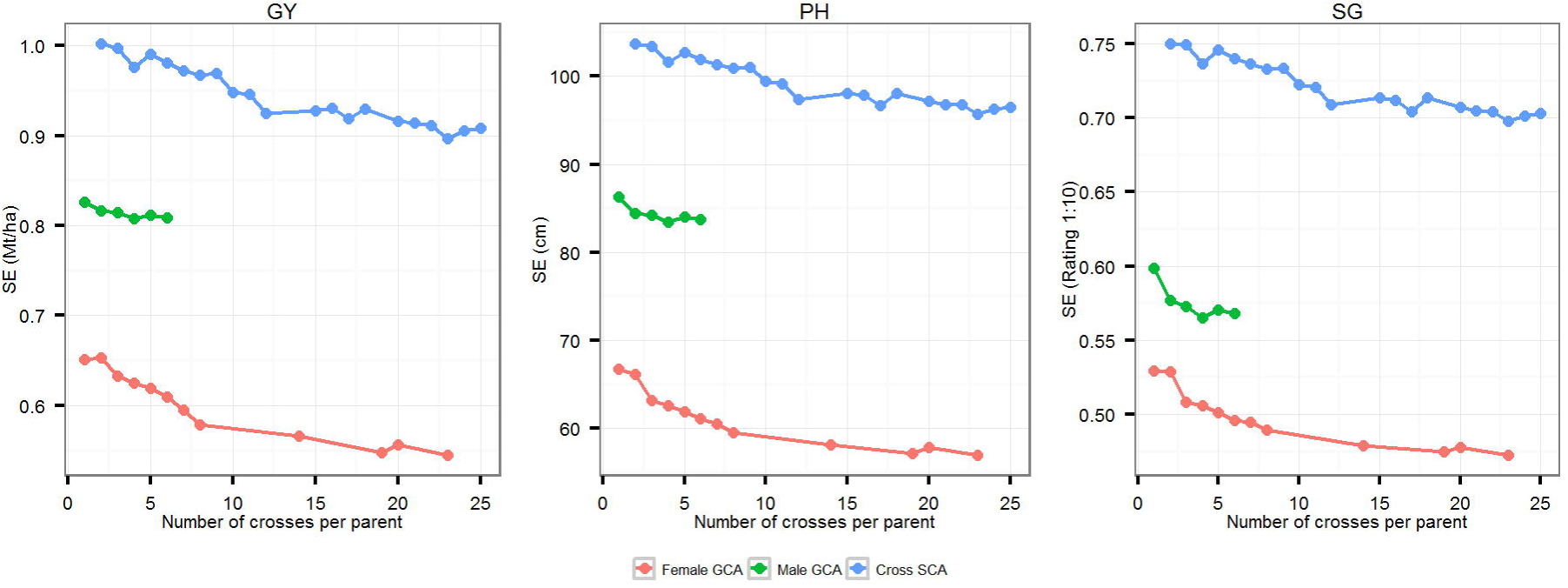
Standard errors of predicted GCA and SCA effects estimated using differing numbers of single crosses per parental inbred progeny.

### Comparison of prediction methods

The published studies on prediction of single cross performance have used covariance among tested and untested crosses (Method 2a & 2b) to predict the performance of untested crosses (Massman *et al.* 2013; Technow *et al.* 2014). In an alternate approach, we used genomic estimated GCA and SCA (Method 1a & 1b) to predict the performance of untested single crosses. The comparison of prediction accuracies showed that the four methods achieved comparable accuracies for predicting T2 and T1F single crosses. For T1M and T0 hybrids, however, Method 1a and 1b provided higher prediction accuracies compared to Method 2a and 2b. Although the two groups of methods use the same information (i.e., additive genomic relationship matrices of inbred parents, dominance relationship matrix of the crosses and tested hybrid data), there are two noteworthy differences between the methods in using this information. First, methods 1a and 1b use the three genomic relationship matrices separately to predict the GCA of each parent and SCA of the cross. Methods 2a and 2b, on the other hand, combine these matrices to calculate the covariance matrices among the tested and untested single crosses. Secondly, in case of method 1a and 1b, all the tested hybrid data is summed by parent and utilized for estimating female and male GCA and cross SCA, while, methods 2a and 2b, use the tested hybrid data as such. The differences in prediction accuracies between two groups of methods for T1M and T0 single crosses can be explained by the above differences in use of the information by these methods. In general, the accuracy of BLUP for untested genotype depends on availability and precise use of information from related tested genotypes and the accuracy of information (or records) on these tested genotypes. The prior part depends on the covariance which is a function of relationship between the tested and untested genotype and genetic variance. When a single population is under consideration, two individuals having high covariance are expected to be more closely related than two individuals having lower covariance. However, when two separate populations are under consideration, two individuals with higher covariance from the first population would not necessarily be more closely related than two individuals with a lower covariance from second population. The reason is that estimate of covariance is population specific as it depends on the genetic variance within a population (Falconer *et al.* 1996) which could differ between populations. As a result, hybrids, where two populations are involved (female population and male population), having higher covariance (as estimated in this and previous studies) may not be more related than two hybrids having lower covariance. Methods 2a and 2b, however, invariably assumes that hybrids having higher covariance are more related than hybrids with lower covariance.

Consider an example of BLUP for T1M hybrids and assume that GCA variance of female is greater than males and compare the two groups of methods in terms of using information from relatives. The methods 1a and 1b, as a result of separately estimating GCA of females and males, will use maximum information from most related tested female to predict the GCA of untested female as GCA variance is constant for female (covariance is simply function of genetic relationship) and covariance between males is not considered in estimating GCA of untested female. Contrastingly, in case of methods 2a and 2b, the information utilized from given tested hybrids depends on covariance between tested and untested hybrid. The hybrid covariance is calculated by summation of covariance between the female parents and covariance between the male parents. For the reasons explained above, the maximum information may not be extracted from most related hybrid because GCA variances of female population and male population are not equal. Specifically, the relationship between female parents, although the male parent is tested for T1M hybrid, is given more weight because GCA variance of females is greater than males. This could result in more use of information from comparatively less related hybrid which affect the prediction accuracy. Now, consider the accuracy of observations on tested genotypes which can also affect the BLUP accuracy. In case of methods 1a and 1b, the GCA of tested parent, which takes into account observations on all tested hybrid where given parent is involved is used for estimating the GCA of untested parent. In case of methods 2a and 2b, the individual observation on hybrid is used for estimating the performance of untested hybrids. The GCA would be more accurate than single observation of tested hybrids as prior is the average of more than one observations. Similarly, the differences in the prediction accuracy of two groups of methods for T0 hybrids can be explained. The similar accuracies of both group of methods for T2 hybrids could also be explained based on above considerations. In predicting the T2 hybrids, as individual has highest relationship with itself, method 1a and 1b are expected to use maximum information from parents of the respective hybrids to estimate their GCA and methods 2a and 2b are also expected to use maximum information from other hybrids of the parents of the respective hybrids. Additionally, more than one closely related hybrids (hybrids involving parents of given T2 hybrid) becomes available for predicting T2 hybrid which leads to accurate determination of the performance of given T2 hybrid by method 2a and 2b as well.

To test this hypothesis for differences in prediction accuracy of two groups of methods, we randomly sampled a balanced (equal females and males) subset of hybrids among 40 females and 40 males. The methods 2a and 2b were modified to correct the discrepancies in weighting of relationship between females and males by using average of GCA variance of females and males. Specifically, for method 2a, the elements of *C*_*ut*_ and off diagonal elements of *C*_*tt*_ were computed as 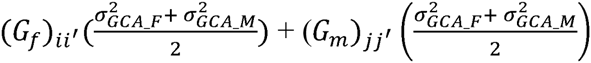. The diagonal elements of *C*_*tt*_ were estimated as 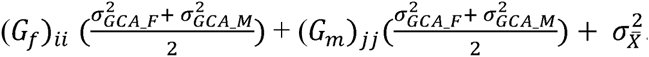. Method 2b was similarly modified. The GY prediction accuracy of four methods for T2, T1F, T1M and T0 hybrid was evaluated using leave-one-individual cross-validations. The modified 2a and 2b methods obtained higher accuracies for T1M and T0 hybrids than original methods 2a and 2b (Table 4). The accuracies were comparable to method 1a and 1b. This confirmed our hypothesis for different prediction accuracy of two groups of methods. For T2 and T1F hybrids, however, the modified 2a and 2b obtained lower accuracies than the original methods. This could also be explained based on our hypothesis. The use of average GCA variance enabled the model to extract information of closer related tested hybrids. However, the amount information extracted depends on variance. The T2 and T1F hybrid involve tested female parent. As GCA variance of females in larger, use of average GCA variance lowered the amount of information extracted due to covariance between female parents. Overall, these results indicate that hybrid covariance based methods, i.e. method 2a and 2b, are inferior in terms of using information from tested hybrids. The combining ability based methods, i.e. methods 1a and 1b, correctly uses genetic relationship and genetic variance by separately estimating female and male GCA. The previous studies on hybrid prediction in maize (Schrag *et al.* 2009; Schrag *et al.* 2010; Technow *et al.* 2014), wheat (Longin *et al.* 2013), sunflower (Reif *et al.* 2013) and triticale (Gowda *et al.* 2013) have reported different estimates of GCA variance between two parental populations of hybrids. This coupled with more accuracy of GCA estimate compared to individual observation on hybrid suggest hybrid prediction based on genomic estimated GCA and SCA is better approach compared to genomic covariance method commonly used.

**Table 4.**
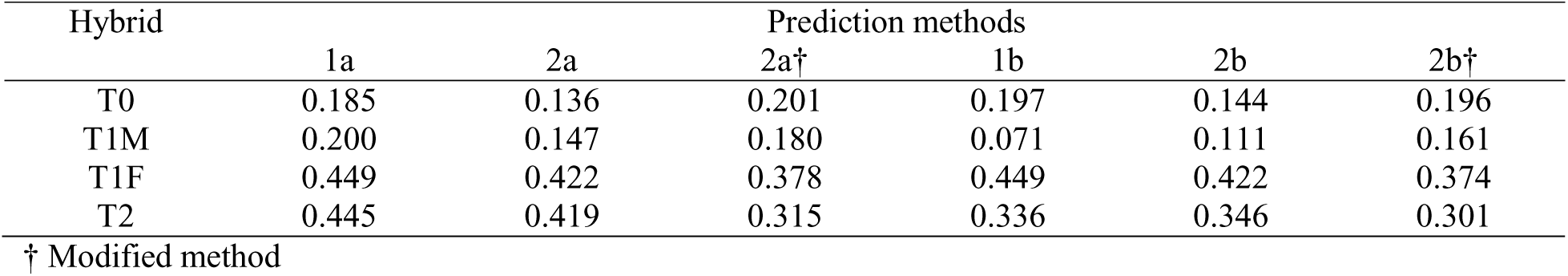
Correlation between observed and predicted grain yield (GY) in a random balanced subset of hybrids for three groups of hybrid prediction methods as evaluated by leave-one-individual-out cross-validation

| Hybrid | Prediction methods |  |  |  |  |  |
| --- | --- | --- | --- | --- | --- | --- |
|  | 1a | 2a | 2a† | 1b | 2b | 2b† |
| T0 | 0.185 | 0.136 | 0.201 | 0.197 | 0.144 | 0.196 |
| T1M | 0.200 | 0.147 | 0.180 | 0.071 | 0.111 | 0.161 |
| T1F | 0.449 | 0.422 | 0.378 | 0.449 | 0.422 | 0.374 |
| T2 | 0.445 | 0.419 | 0.315 | 0.336 | 0.346 | 0.301 |

† Modified method

The prediction accuracies for T2, T1, and T0 hybrids reported by (Technow *et al.* 2014) and T2 and T1 hybrids reported by (Massman *et al.* 2013) are higher than corresponding accuracies observed in the present study. This discrepancy is likely due to the differences in population and family structure between the present study and those previously reported. Let’s consider two prediction scenarios, one from these previous studies and one in our study. Massman *et al.* (2013) and Technow *et al.* (2014) have used single crosses made among diverse set of established inbred parents. These inbred parents are likely to belong distinct groups based on their hybrid performances. As Windhausen *et al.* (2012) reported, the prediction accuracy under such scenario results mostly from differences in mean performances between groups and less from genetic relationship between training and validation set because large amount variation happen to be between groups compared to within groups. In our case, larger genetic variation was within families because there were many inbred progenies from each bi-parental family and there was one grandparent common between the families. Therefore, hybrid prediction accuracy was mostly resulted from genetic relationship and less from differences in mean performances between groups. However, the closer genetic relationship between training and validation set generated due to the common grandparent made challenging to distinguish the hybrid performances among closely related inbred progenies. In addition, the average number of hybrid combinations per parental line was higher in these studies which significantly increases the hybrid prediction accuracy as evidenced from higher prediction accuracy of T1F crosses compared to T1M crosses

### The benefit of modeling SCA

We observed an increase in prediction accuracy for GY by modeling and estimating SCA effects and subsequently summing GCA and SCA effects. The increase in accuracy by modeling SCA was highest for T0 hybrids followed by T1 and T2 hybrids. This result suggests that modeling SCA can be more beneficial for hybrids with untested parents compared to hybrids with one or two tested parents. These findings could possibly be explained by a small number of hybrid combinations per parental line. If parents are tested in a small number of hybrid combinations, as in the present study, their GCA effect predictions could capture a significant portion of the SCA effect as well. The increase in prediction accuracy achieved by adding SCA would clearly depend on the magnitude of the SCA bias of the predicted GCA effect. We do not have the ability to estimate this bias, but in our study the ratio of SCA vs. GCA variance was small for all traits (Table 3). When a parent has no performance data in hybrid combination, its GCA is predicted based on all tested relatives, resulting in a predicted GCA effect less biased by SCA. Hence, SCA is expected to improve the predictions for hybrids with untested parents. The highest increase in prediction accuracy by modeling SCA covariance was achieved for GY followed by PH, and SG. This trend can be explained by a higher proportion of in total genetic variance for GY compared to PH and SG (Table 3).

When predicting the GY of hybrids from the novel family (i.e. leave-one-family-out cross-validations), the accuracy was similar or slightly lower when both GCA and SCA used in comparison to using only GCA for prediction. However, when hybrids from novel family added in the training set (i.e. leave-one-individual-out cross-validation), the prediction accuracy generally increased with method 1b. When no information was available from hybrids within a novel family, SCA is determined from hybrid combinations of relatives in other families, which are expected to be less accurate since SCA depends on specific parental combinations. Alternatively, when some hybrids from the same family are present in training set, they are used in SCA estimation which is expected to be more accurate because of closer relatedness among hybrid combinations in this case. Overall, this suggest that using SCA in addition to GCA for hybrid prediction is beneficial when closer related hybrids (i.e. same single-cross family) are available and GCA estimates are less biased.

The previous experimental studies in maize (Bernardo. 1994), wheat (Zhao *et al.* 2013), triticale (Gowda *et al.* 2013) and sunflower (Reif *et al.* 2013) reported a small decrease in prediction accuracy by modelling SCA effect in addition to GCA effects. These studies have used diverse set of inbreds. As we observed in predicting hybrids from novel family, this decrease in prediction accuracy can be attributed to inability to accurately predict SCA from distantly related hybrid combinations.

### Efficiency of early-stage hybrid prediction

Although we don’t have topcross data, the comparisons of “early-stage hybrid prediction” suggested in this study with current practice of “topcross based initial selections” could be made. There are total 217 inbred progenies used in this study. Single tester based selection would need 217 test crosses to be made for first screening of inbreds. This is followed by additional test crosses of inbreds selected from first screening with more testers. Now, our results are based on training set size of 250. Therefore, number of crosses required to be made can be assumed comparable. Two important differences, however, can easily be seen between the two schemes. First, the breeding cycle of hybrid development is short in “early-stage hybrid prediction” compared to “topcross based initial selections”. Secondly, in “early-stage hybrid prediction”, actual GCA of inbreds and SCA of all crosses are estimated and, subsequently, used to determine the genetic value of all potential hybrid combinations (i.e. 7866). The prediction accuracy for unobserved hybrid combinations is expected to be similar to T2 scenario as each parent of hybrid is tested in hybrid combinations (Zhao *et al.* 2015). Hence, superior single crosses are identified with high certainty. In case of “topcross based initial selections”, early selections among available inbreds are based on test cross value and only hybrid combinations among selected inbreds are evaluated in later stages. There is good possibility of losing some superior hybrid combinations because SCA is important component of hybrid performance as indicated by significant SCA variance for all traits in the present study (Table 3). Overall, this suggest that genomic prediction of the performance of single crosses made using random progenies from the early stages of the breeding pipeline holds great potential to re-design hybrid breeding and increase its efficiency.

### Prospects for optimization of genomic hybrid prediction

Pedigree selection and frequent use of successful parents creates a family structure within typical hybrid maize breeding programs consisting of inter-connected bi-parental families. This family structure should be leveraged to optimize hybrid genomic prediction training sets. If a common training set useful across large segments of the breeding germplasm could be designed, significant resources could be saved. The results from this study not only show that prediction of single cross performance holds great potential, it also demonstrates its potential during the very early stages of a breeding program where breeding populations are comprised of many random progenies from large bi-parental families. Along these lines, we demonstrated that hybrid genomic prediction methods even hold potential for separating single crosses from a common family background (Figure 4). The prediction accuracy of single crosses across families was accurate and the addition of single crosses from the same family to the training set only minimally improved accuracy. This finding suggests that training sets can be formed and used to predict related families not represented in the training set. Further study of the optimization of larger training sets through leveraging family structure will likely improve accuracies far beyond those measured herein.

---

**Supplementary Figure 1.**
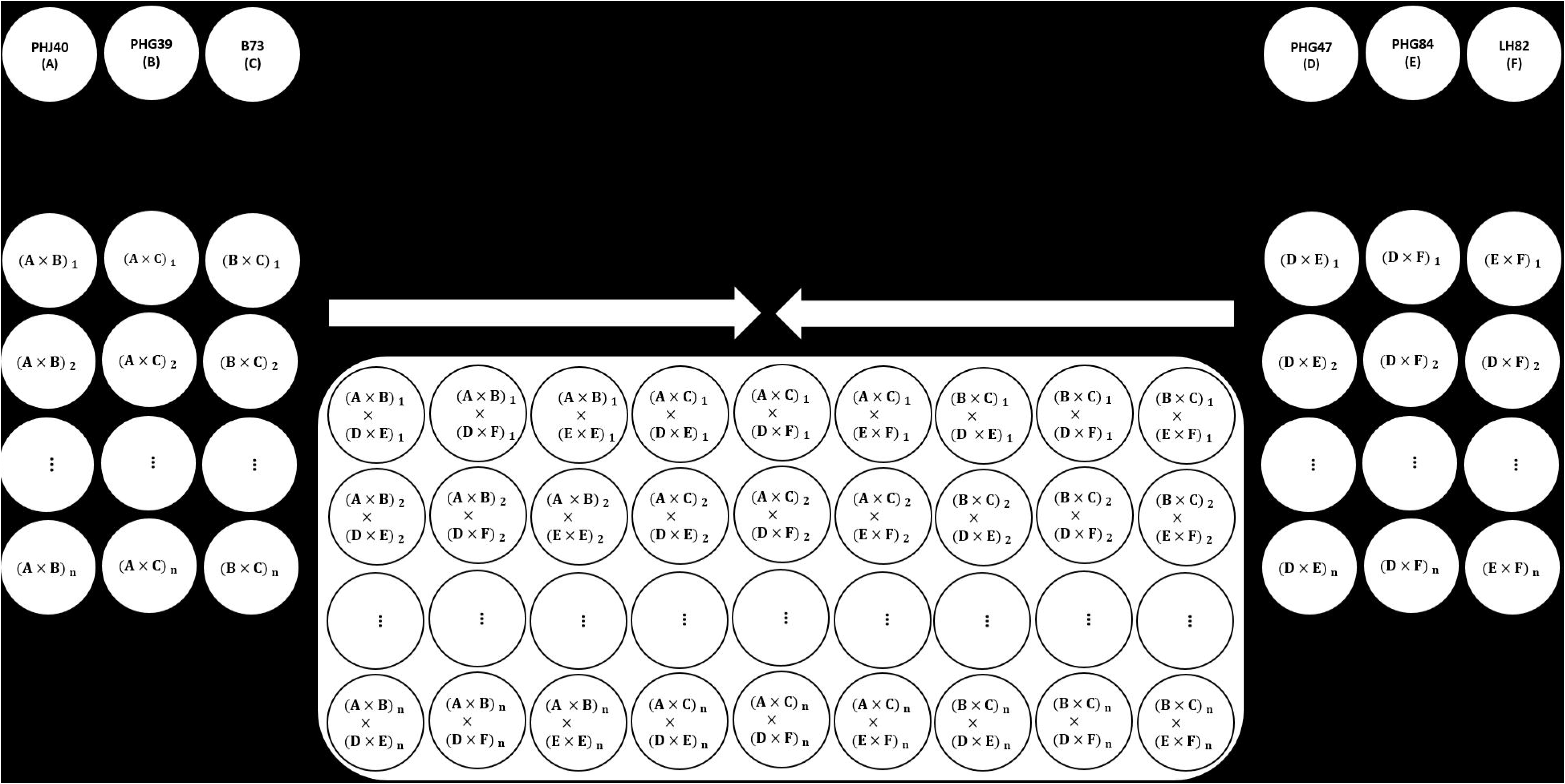
The connected bi-parental population structure. Six inbreds were selected as parents for the connected bi-parental population based on the results of the plant density tolerance survey by Mansfield and Mumm. (2014). RILs and DH lines were available for each of the three possible crosses within the heterotic groups. These RILs and DH lines from the SSS and NSS families were crossed to maximize the number of RIL and DH parents used in the creation of the population, as well as maintain the balance of individuals in each of the nine single-cross families.

**Supplementary Figure 2.**
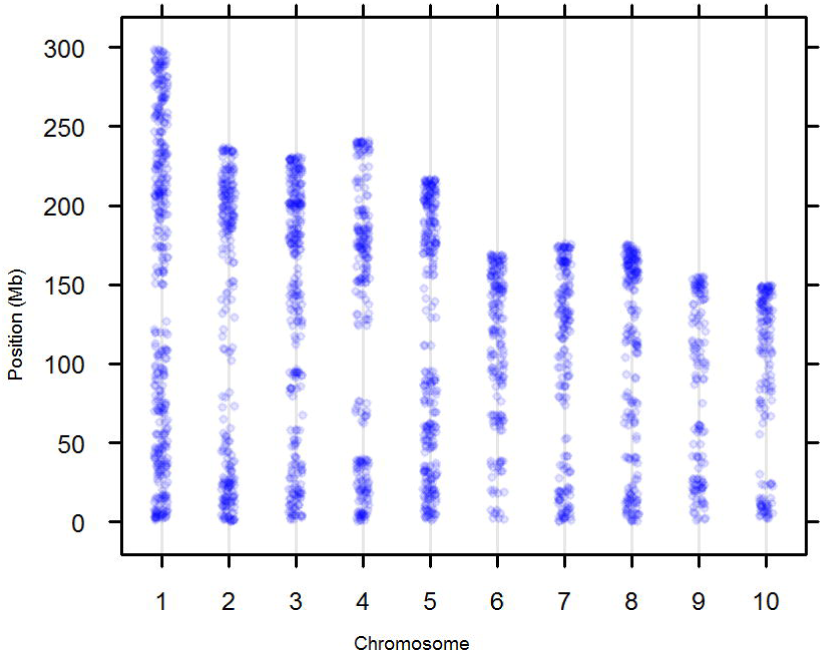
Distribution of 2296 single nucleotide polymorphisms scored using genotyping by sequencing on the ten chromosomes of maize

**Table S1.**
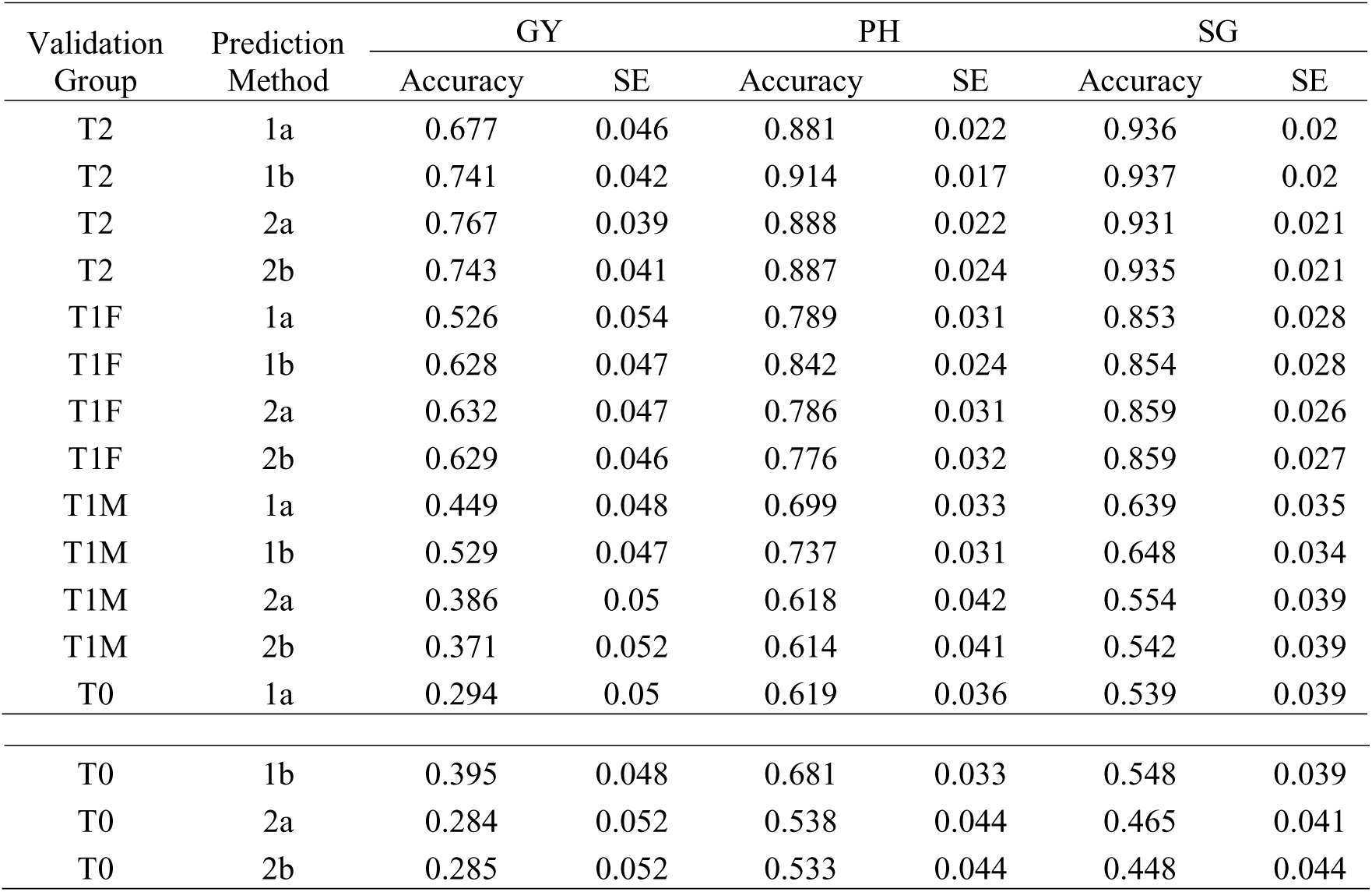
Prediction accuracy for T2, T1F, T1M and T0 hybrids for traits GY, PH and SG obtained using the four methods 1a (Parent GCA), 1b (Parent GCA plus single-cross SCA), 2a (Additive genetic covariance among single crosses) and 2b (Additive plus dominance covariance among single crosses) as evaluated with training set of 250 and leave-one-individual-out cross-validation

| Validation Group | Prediction Method | GY |  | PH |  | SG |  |
| --- | --- | --- | --- | --- | --- | --- | --- |
|  |  | Accuracy | SE | Accuracy | SE | Accuracy | SE |
| T2 | 1a | 0.677 | 0.046 | 0.881 | 0.022 | 0.936 | 0.02 |
| T2 | 1b | 0.741 | 0.042 | 0.914 | 0.017 | 0.937 | 0.02 |
| T2 | 2a | 0.767 | 0.039 | 0.888 | 0.022 | 0.931 | 0.021 |
| T2 | 2b | 0.743 | 0.041 | 0.887 | 0.024 | 0.935 | 0.021 |
| T1F | 1a | 0.526 | 0.054 | 0.789 | 0.031 | 0.853 | 0.028 |
| T1F | 1b | 0.628 | 0.047 | 0.842 | 0.024 | 0.854 | 0.028 |
| T1F | 2a | 0.632 | 0.047 | 0.786 | 0.031 | 0.859 | 0.026 |
| T1F | 2b | 0.629 | 0.046 | 0.776 | 0.032 | 0.859 | 0.027 |
| T1M | 1a | 0.449 | 0.048 | 0.699 | 0.033 | 0.639 | 0.035 |
| T1M | 1b | 0.529 | 0.047 | 0.737 | 0.031 | 0.648 | 0.034 |
| T1M | 2a | 0.386 | 0.05 | 0.618 | 0.042 | 0.554 | 0.039 |
| T1M | 2b | 0.371 | 0.052 | 0.614 | 0.041 | 0.542 | 0.039 |
| T0 | 1a | 0.294 | 0.05 | 0.619 | 0.036 | 0.539 | 0.039 |
| T0 | 1b | 0.395 | 0.048 | 0.681 | 0.033 | 0.548 | 0.039 |
| T0 | 2a | 0.284 | 0.052 | 0.538 | 0.044 | 0.465 | 0.041 |
| T0 | 2b | 0.285 | 0.052 | 0.533 | 0.044 | 0.448 | 0.044 |

**Table S2.** Accuracy of predicting the hybrids from novel family obtained using the methods 1a (Parent GCA), 1b (Parent GCA plus single-cross SCA) as evaluated with training set of 250 and leave-one-individual-out and leave-one-family-out cross-validations for traits GY, PH and SG.

| Family | GY |  |  |  | PH |  |  |  | SG |  |  |  |
| --- | --- | --- | --- | --- | --- | --- | --- | --- | --- | --- | --- | --- |
|  | Leave one individual out |  | Leave one family out |  | Leave one individual out |  | Leave one family out |  | Leave one individual out |  | Leave one family out |  |
|  | 1a | 1b | 1a | 1b | 1a | 1b | 1a | 1b | 1a | 1b | 1a | 1b |
| f1 | 0.528 | 0.671 | 0.79 | 0.647 | 0.856 | 0.854 | 0.854 | 0.811 | 0.793 | 0.79 | 0.790 | 0.854 |
| f2 | 0.847 | 0.840 | 0.874 | 0.715 | 0.869 | 0.876 | 0.876 | 0.842 | 0.857 | 0.87 | 0.870 | 0.807 |
| f3 | 0.396 | 0.261 | 0.545 | 0.49 | 0.883 | 0.879 | 0.879 | 0.929 | 0.828 | 0.816 | 0.816 | 0.949 |
| f4 | 0.620 | 0.714 | 0.544 | 0.565 | 0.834 | 0.861 | 0.861 | 0.8 | 0.784 | 0.775 | 0.775 | 0.712 |
| f5 | 0.702 | 0.600 | 0.771 | 0.717 | 0.819 | 0.83 | 0.830 | 0.767 | 0.889 | 0.872 | 0.872 | 0.923 |
| f6 | 0.525 | 0.550 | 0.487 | 0.556 | 0.772 | 0.802 | 0.802 | 0.421 | 0.580 | 0.575 | 0.575 | 0.453 |

## LITERATURE CITED

Albrecht, T., H. -. Auinger, V. Wimmer, J. O. Ogutu, C. Knaak et al, 2014 Genome-based prediction of maize hybrid performance across genetic groups, testers, locations, and years. Theor. Appl. Genet. 127: 1375–1386.

Albrecht, T., V. Wimmer, H. Auinger, M. Erbe, C. Knaak et al, 2011 Genome-based prediction of testcross values in maize. Theor. Appl. Genet. 123: 339–350.

Bernardo, R., 1992 Relationship between single-cross performance and molecular marker heterozygosity. Theor. Appl. Genet. 83: 628–634.

Bernardo, R., 2002 Breeding for Quantitative Traits in Plants. Stemma Press Woodbury.

Bernardo, R., 1996a Best linear unbiased prediction of maize single-cross performance. Crop Sci. 36: 50–56.

Bernardo, R., 1996b Best linear unbiased prediction of the performance of crosses between untested maize inbreds. Crop Sci. 36: 872–876.

Bernardo, R., 1994 Prediction of maize single-cross performance using RFLPs and information from related hybrids. Crop Sci. 34: 20–25.

Butler, D., B. R. Cullis, A. Gilmour and B. Gogel, 2009 ASReml-R reference manual. The State of Queensland, Department of Primary Industries and Fisheries, Brisbane.

Canty, A. J., 2014 EEF. profile 47. Package ‘boot’ 47.

Charcosset, A., M. Lefort-Buson and A. Gallais, 1991 Relationship between heterosis and heterozygosity at marker loci: A theoretical computation. Theoret. Appl. Genetics 81: 571–575.

Comstock, R. E., and H. F. Robinson, 1948 The components of genetic variance in populations of biparental progenies and their use in estimating the average degree of dominance. Biometrics 4: 254–266.

de los Campos, G., J. M. Hickey, R. Pong-Wong, H. D. Daetwyler and M. P. L. Calus, 2013 Whole-genome regression and prediction methods applied to plant and animal breeding. Genetics 193: 327–345.

Dekkers, J., 2007 Marker-assisted selection for commercial crossbred performance. J. Anim. Sci. 85: 2104–2114.

Elshire, R. J., J. C. Glaubitz, Q. Sun, J. A. Poland, K. Kawamoto et al, 2011 A robust, simple genotyping-by-sequencing (GBS) approach for high diversity species. PLoS One 6: e19379.

Falconer, D. S., T. F. Mackay and R. Frankham, 1996 Introduction to quantitative genetics (4th edn). Trends in Genetics 12: 280.

Fehr, W., 1987 Principles of cultivar development: Theorey and technique.

Glaubitz, J. C., T. M. Casstevens, F. Lu, J. Harriman, R. J. Elshire et al, 2014 TASSEL-GBS: A high capacity genotyping by sequencing analysis pipeline. PLoS One 9: e90346.

Gowda, M., Y. Zhao, H. P. Maurer, E. A. Weissmann, T. Würschum et al, 2013 Best linear unbiased prediction of triticale hybrid performance. Euphytica 191: 223–230.

Hallauer, A. R., 1977 Relation between inbred and hybrid traits in maize. Crop Sci. 17: 703–706.

Hallauer, R., W. Russell and K. Lamkey, 1988 Corn breeding. Corn and Corn Improvement 463–564.

He, J., X. Zhao, A. Laroche, Z. Lu, H. Liu et al, 2014 Genotyping-by-sequencing (GBS), an ultimate marker-assisted selection (MAS) tool to accelerate plant breeding. Frontiers in plant science 5:.

Heffner, E. L., M. E. Sorrells and J. Jannink, 2009 Genomic selection for crop improvement. Crop Sci. 49: 1–12.

Holland, J. B., W. E. Nyquist and C. T. Cervantes-Martínez, 2003 Estimating and interpreting heritability for plant breeding: An update. Plant Breed. Rev. 22: 9–112.

Jacobson, A., L. Lian, S. Zhong and R. Bernardo, 2014 General combining ability model for genomewide selection in a biparental cross. Crop Sci. 54: 895–905.

Jenkins, M. T., and A. M. Brunson, 1932 Methods of testing inbred lines of maize in crossbre combinations. J.Am.Soc.Agron 24: 523–530.

Lee, E., M. Ash and B. Good, 2007 Re-examining the relationship between degree of relatedness, genetic effects, and heterosis in maize. Crop Sci. 47: 629–635.

Lin, Z., B. Hayes and H. Daetwyler, 2014 Genomic selection in crops, trees and forages: A review. Crop and Pasture Science 65: 1177–1191.

Longin, C. F. H., M. Gowda, J. Mühleisen, E. Ebmeyer, E. Kazman et al, 2013 Hybrid wheat: Quantitative genetic parameters and consequences for the design of breeding programs. Theor. Appl. Genet. 126: 2791–2801.

Love, H., and B. Wentz, 1914 Correlations between ear characters and yield in corn. Laboratory Papers.V.1 3:.

Mansfield, B. D., and R. H. Mumm, 2014 Survey of plant density tolerance in US maize germplasm. Crop Sci. 54: 157–173.

Massman, J. M., A. Gordillo, R. E. Lorenzana and R. Bernardo, 2013 Genomewide predictions from maize single-cross data. Theor. Appl. Genet. 126: 13–22.

Melchinger, A., 1999 Genetic diversity and heterosis. The genetics and exploitation of heterosis in crops 10:.

Meuwissen, T. H., B. J. Hayes and M. E. Goddard, 2001 Prediction of total genetic value using genome-wide dense marker maps. Genetics 157: 1819–1829.

Patterson, H., and E. Williams, 1976 A new class of resolvable incomplete block designs. Biometrika 63: 83–92.

Reif, J. C., Y. Zhao, T. Würschum, M. Gowda and V. Hahn, 2013 Genomic prediction of sunflower hybrid performance. Plant Breeding 132: 107–114.

Riedelsheimer, C., A. Czedik-Eysenberg, C. Grieder, J. Lisec, F. Technow et al, 2012 Genomic and metabolic prediction of complex heterotic traits in hybrid maize. Nat. Genet. 44: 217–220.

Schrag, T., M. Frish, B. Dhillon and A. Melchinger, 2009 Marker-based prediction of hybrid performance in maize single-crosses involving doubled haploids. Maydica 54: 353.

Schrag, T., A. Melchinger, A. Sørensen and M. Frisch, 2006 Prediction of single-cross hybrid performance for grain yield and grain dry matter content in maize using AFLP markers associated with QTL. Theor. Appl. Genet. 113: 1037–1047.

Schrag, T. A., H. P. Maurer, A. E. Melchinger, H. Piepho, J. Peleman et al, 2007 Prediction of single-cross hybrid performance in maize using haplotype blocks associated with QTL for grain yield. Theor. Appl. Genet. 114: 1345–1355.

Schrag, T. A., J. Möhring, A. E. Melchinger, B. Kusterer, B. S. Dhillon et al, 2010 Prediction of hybrid performance in maize using molecular markers and joint analyses of hybrids and parental inbreds. Theor. Appl. Genet. 120: 451–461.

Shull, G. H., 1909 A pure-line method in corn breeding. J. Hered. 51–58.

Smith, C. W., and J. Betrán, 2004 Corn: Origin, History, Technology, and Production. John Wiley & Sons.

Smith, O., 1986 Covariance between line per se and testcross performance. Crop Sci. 26: 540–543.

Technow, F., C. Riedelsheimer, T. A. Schrag and A. E. Melchinger, 2012 Genomic prediction of hybrid performance in maize with models incorporating dominance and population specific marker effects. Theor. Appl. Genet. 125: 1181–1194.

Technow, F., T. A. Schrag, W. Schipprack, E. Bauer, H. Simianer et al, 2014 Genome properties and prospects of genomic prediction of hybrid performance in a breeding program of maize. Genetics 197: 1343–1355.

VanRaden, P., 2008 Efficient methods to compute genomic predictions. J. Dairy Sci. 91: 4414–4423.

Vuylsteke, M., M. Kuiper and P. Stam, 2000 Chromosomal regions involved in hybrid performance and heterosis: Their AFLP®-based identification and practical use in prediction models. Heredity 85: 208–218.

Windhausen, V. S., G. N. Atlin, J. M. Hickey, J. Crossa, J. L. Jannink et al, 2012 Effectiveness of genomic prediction of maize hybrid performance in different breeding populations and environments. G3 (Bethesda) 2: 1427–1436.

Zhao, Y., J. Zeng, R. Fernando and J. C. Reif, 2013 Genomic prediction of hybrid wheat performance. Crop Sci. 53: 802–810.

Zhao, Y., Z. Li, G. Liu, Y. Jiang, H. P. Maurer et al, 2015 Genome-based establishment of a high-yielding heterotic pattern for hybrid wheat breeding. Proc. Natl. Acad. Sci. U. S. A. 112: 15624–15629.

